## SupplementalFigures_Tables for "*Cryptococcus neoformans* Slu7 ensures nuclear positioning during mitotic progression through RNA splicing"

**Supplementary figures and tables**

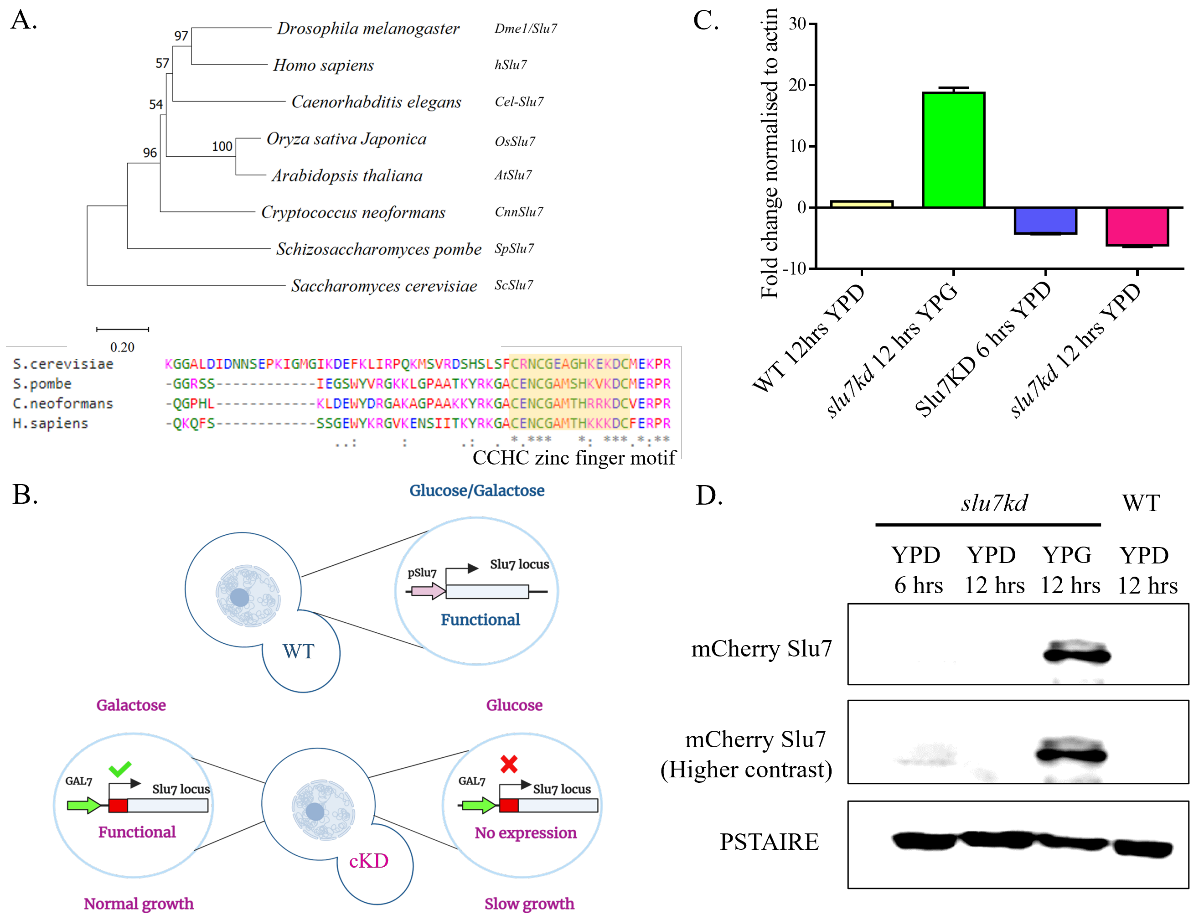


**Supplementary Figure 1 : CnSlu7 is evolutionary close to higher eukaryotes than the other fungal orthologs and its depletion results in slow growth A**. Phylogenetic tree conservation of Slu7 protein among eukaryotes. The tree was generated using MEGA 11 with neighbor-joining method and the following parameters: Poisson correction, pairwise deletion, and bootstrap (1,000 replicates). The zinc finger motif conserved in fungal species and humans. **B.** Schematic representation of Slu7 knockdown strain. **C.** Quantification of RNA depletion in *slu7kd* and wildtype cells after the shift into non-permissive media for 6 hours and 12 hours by qRT PCR. **D.** Western blot to detect mCherry tagged Slu7 protein in *slu7kd* grown in non-permissive media for 6 hours and 12 hours at 30°C. The blot was probed with anti-mCherry as described in the Materials and Method section. pSTAIRE was used as loading control. Knockdown strain grown in permissive media for 12 hours was used as positive control.

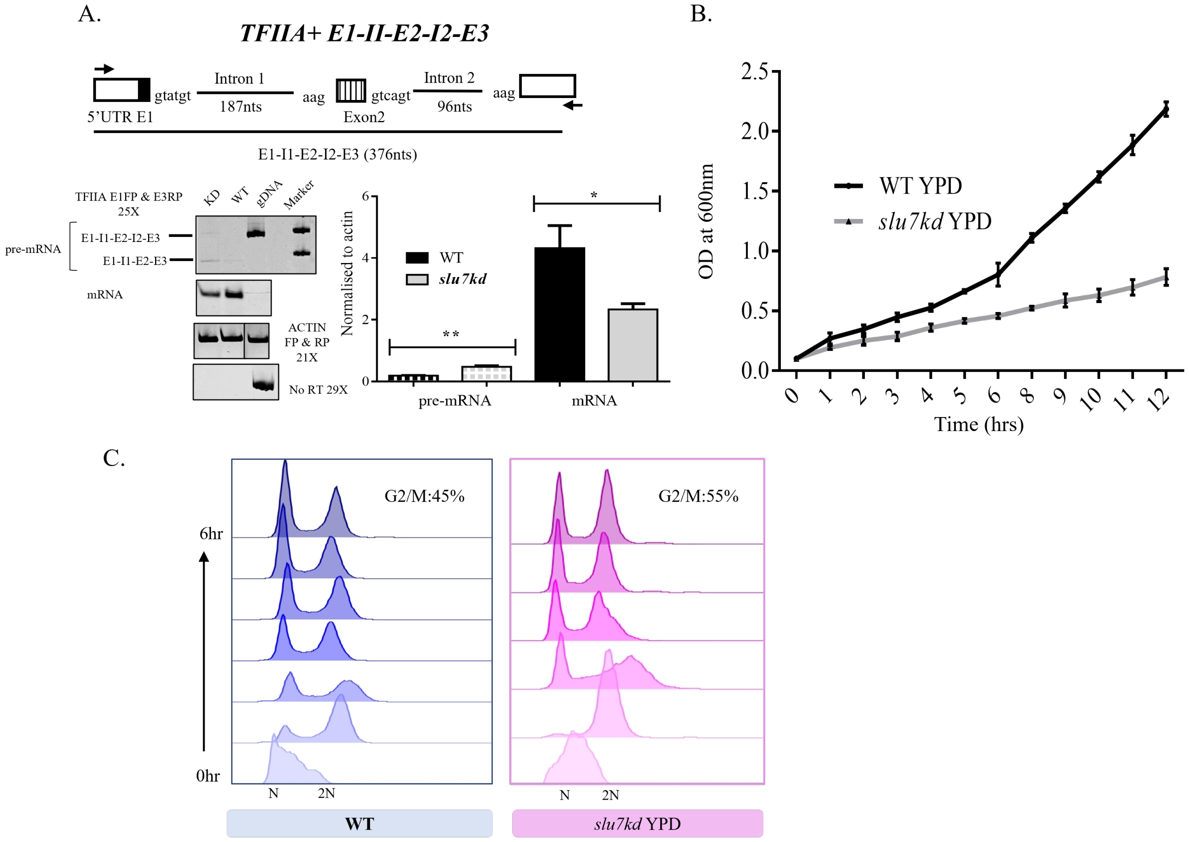


**Supplementary Figure 2: Conditional knockdown of Slu7 shows splicing defect of *TFIIA intron 1* and slower progression in mitosis A.** Splicing defect of TFIIA intron 1 in *slu7kd* and wildtype grown in YPD for 12 hours. **B.** Growth profile of broth cultures of wildtype and *slu7kd* grown in non-permissive media. The data represent mean ± SD for three independent biological replicates. **C.** Flow cytometry analysis of synchronous wildtype and Slu7 conditional knockdown after release into non-permissive media. The percentage at the top represents the % of cells in the G2/M phase at the end of 6 hours, N = 3.

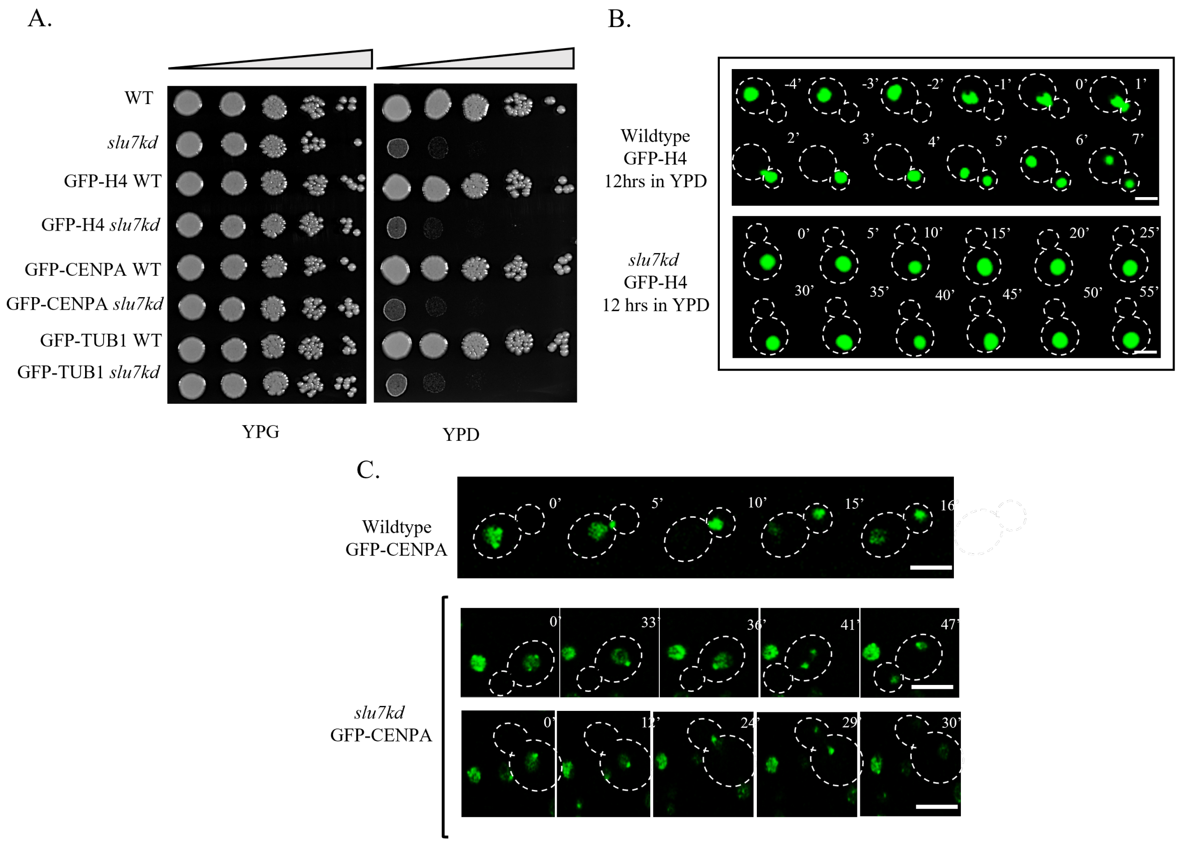


**Supplementary Figure 3 : Dynamics of nucleus and kinetochore patterning in *slu7kd* A.** Serial 10-fold dilution of 2 X $10^5$ cells from wildtype, *slu7kd* strain, each of which had marked reporters for monitoring mitosis. These strains with GFP-H4, GFP-TUB1, and GFP-CENPA reporters were spotted on non-permissive media and monitored for growth at 30°C for 5 days. **B.** Time-lapse snapshots of *slu7kd* and wildtype cells with GFP-H4 reporter to visualize nuclear dynamics after growth in non-permissive media for 12 hours. T = 0 was taken when the nucleus enters the neck region in the wildtype panel. In the knockdown panel, the timestamps are mentioned right from the start of the imaging. Bar, 5µm. **C.** Time-lapse snapshots of *slu7kd* and wildtype cells with GFP-CENPA reporter to visualize the kinetochore dynamics after growth in non-permissive media for 6 hours. T = 0 represents the start of the live imaging. Bar, 5µm.

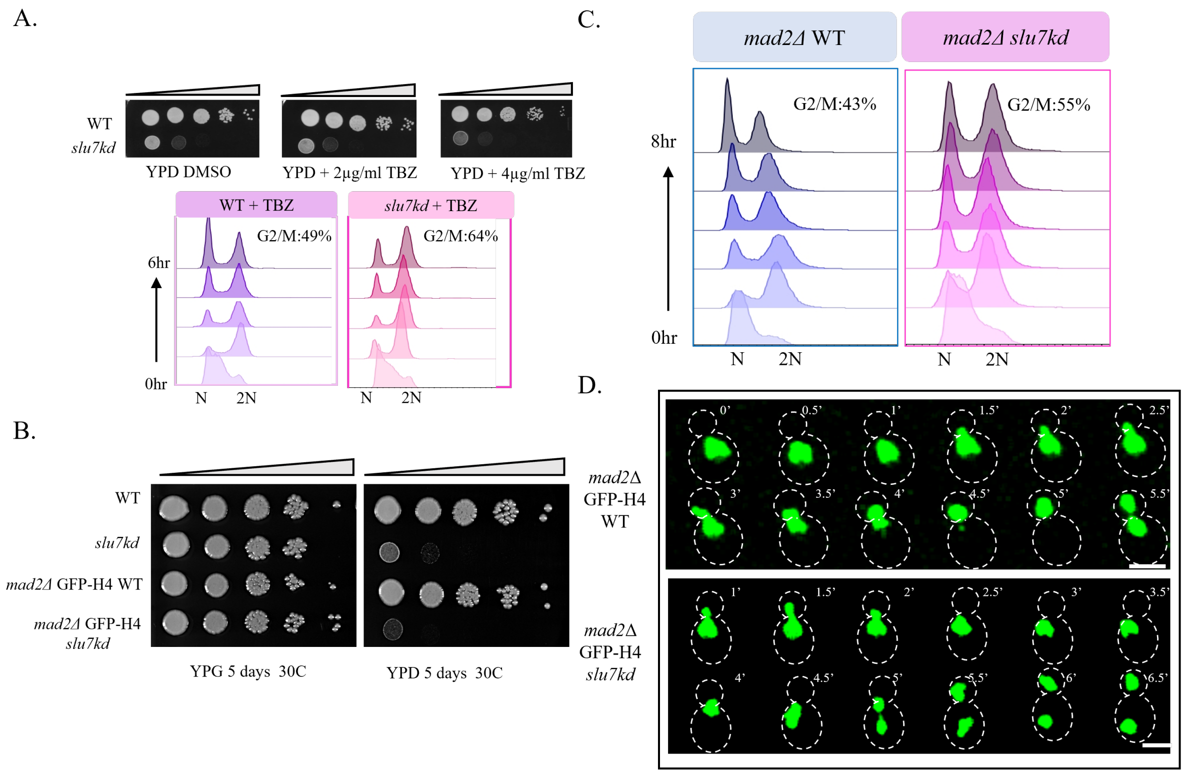


**Supplementary Figure 4 : Slu7 mediated defects are not under the surveillance of Spindle Assembly Checkpoint (SAC) A.** Serial 10-fold dilution of 2 X $10^5$ cells from wildtype and *slu7kd* spotted on non-permissive media containing 2µg/ml and 4µg/ml thiabendazole and monitored for growth at 30°C for 5 days. Flow cytometry analysis of cells from wildtype and *slu7kd* strain withdrawn at various time points after inoculation of HU- synchronised cells into non-permissive media containing 4µg/ml thiabendazole. The percentage figures given at the top represents the % of cells in the G2/M phase at the end of 6 hours, N = 3. **B.** Serial 10-fold dilution of 2 X $10^5$ cells from wildtype and *slu7kd* in the background of *mad2Δ* GFP-H4 spotted on non-permissive media and monitored for growth at 30°C for 5 days. **C.** Flow cytometry analysis of cells from wildtype and *slu7kd* strain in the background of *mad2Δ* GFP-H4 withdrawn at various time points after inoculation of HU- synchronised cells into non-permissive media. The percentage figures given at the top represents the % of cells in the G2/M phase at the end of 8 hours, N = 3. **D.** Time-lapse snapshots of *slu7kd* and wildtype cells in the background of *mad2Δ* GFP-H4 to visualize nuclear dynamics after growth in non-permissive media for 6 hours. T = 0 was taken when the nucleus enters the neck region between the mother and daughter cell. Bar, 5µm.

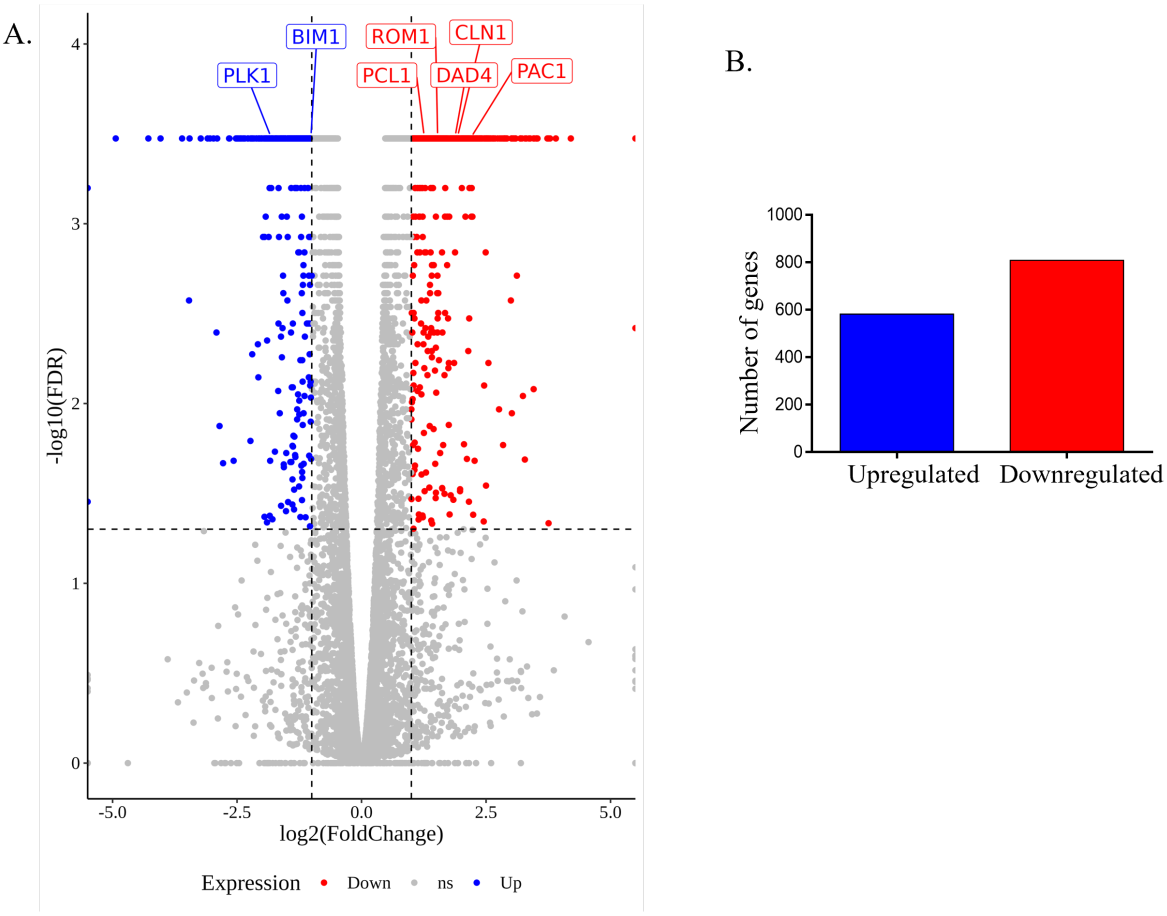


**Supplementary Figure 5 : Slu7 knockdown leads to deregulation of transcripts involved various cellular functions A.** Volcano plot representing the differential gene expression between *slu7kd* and Wildtype. **B.** The bar chart represents the upregulated and downregulated genes in Slu7 knockdown compared to wildtype.

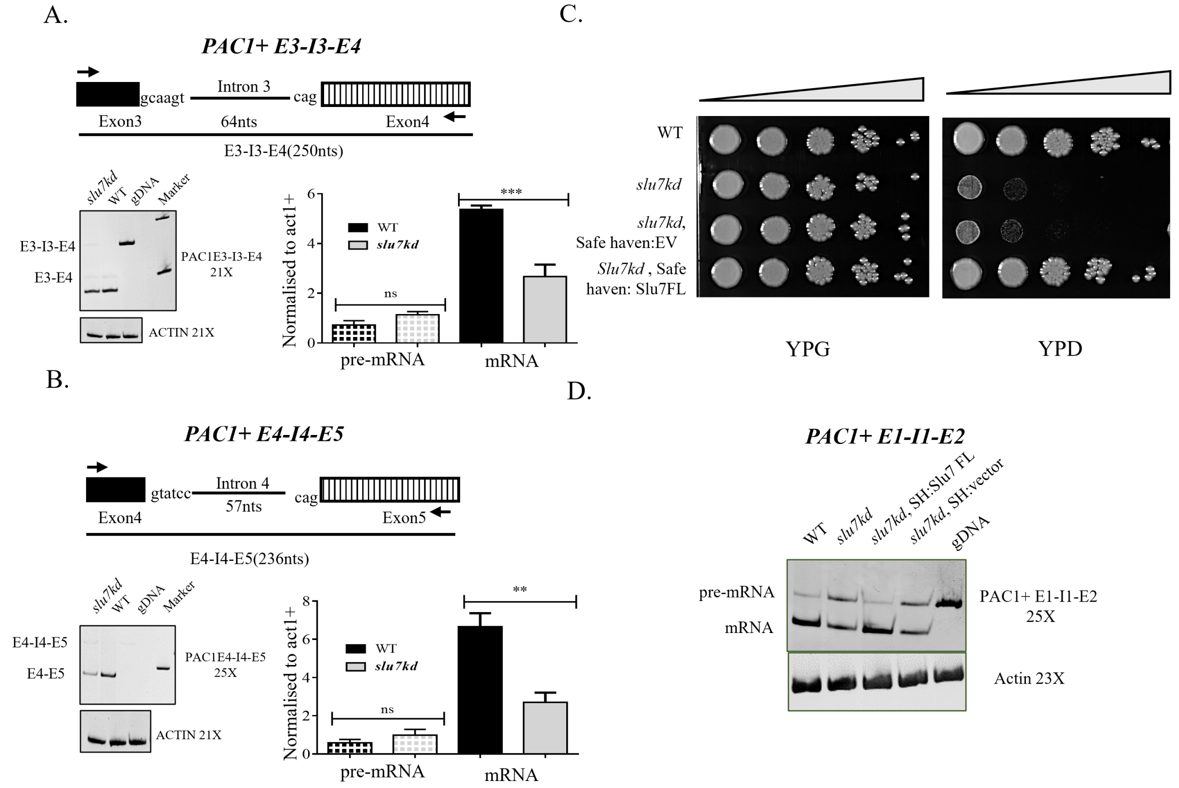


**Supplementary Figure 6 : Intron 3 and intron 4 of PAC1 transcript are not dependent on Slu7 for splicing.**

Schematic representations show each intron together with its flanking exons. Intron length is given within brackets. RNA from WT and *slu7kd* cells grown at 30 °C for 12 hours was taken for limiting cycle, semi-quantitative RT-PCR using the flanking exonic primers. For each intron, the pre-mRNA (P) or mRNA (M) levels were normalized to that of the act1+ (A) mRNA. The normalized pre-mRNA or mRNA levels are plotted. The data represent mean ± SD for three independent biological replicates. p values were determined by unpaired Student's t-test. ns, non-significant change with $p > 0.05$. **A.** The splicing status of the PAC1+ intron 3 in wildtype and *slu7kd*. **B.** The splicing status of the PAC1+ intron 4 in wildtype and *slu7kd*. **C.** Serial 10 dilutions starting from $2 \times 10^5$ cells of wildtype, Slu7 conditional knockdown, and two transformants expressing Slu7FL from safe haven locus in the background of *slu7kd* were spotted on non-permissive media. The image was obtained after incubating the plates at 30°C for 5 days. **D.** The splicing status of the PAC1+ intron 1 in wildtype, *slu7kd*, and *slu7kd* expressing intronless PAC1 from safe haven loci. RNA from WT, Slu7 KD, and Slu7 KD cells expressing intronless PAC1 grown at 30 °C for 12 hours was taken for limiting cycle, semi-quantitative RT-PCR using the flanking exonic primers. The experiment was done in three independent biological replicates.

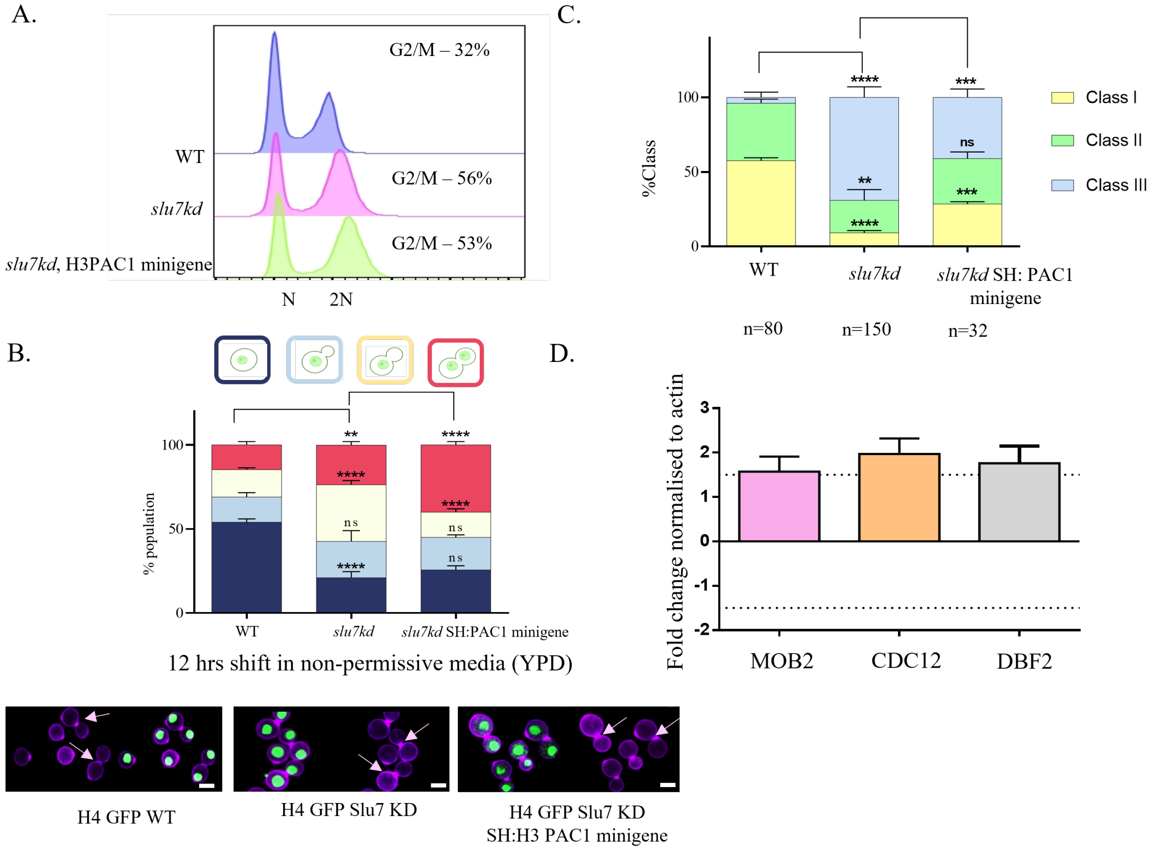


**Supplementary Figure 7 : Slu7KD with intronless PAC1 arrest post nuclear division in late anaphase / cytokinesis A.** Flow cytometry analysis of cells from wildtype, *slu7kd* and *slu7kd* expressing PAC1 minigene grown in non-permissive media (YPD) for 12 hours. **B.** The percentage of cells at various phases of the cell cycle, based on bud and nuclear position was measured using *slu7kd* (n = 100), wildtype (n =100) and *slu7kd* SH::PAC1 minigene (n =100) was measured in fixed cells (4% paraformaldehyde) after growth in YPD for 12hrs, respectively. The data represent mean ± SD for three independent biological replicates. One-way ANOVA test followed by Turkey's multiple comparison test was used to calculate the statistical significance of differences between the population (the p values show the difference compared to the wildtype vs *slu7kd*, *slu7kd* vs *slu7kd* SH:PAC1 minigene). Snapshots of wildtype GFP-H4 WT, *slu7kd* GFP-H4, and *slu7kd* with H3:PAC1intronless minigene GFP-H4 cells stained with calcofluor white to visualize the cell wall. Bar, 5µm. **C.** Localization of Dyn1 in the wildtype, *slu7kd* and *slu7kd* with PAC1 minigene cells at different stages expressing Dyn1-3xGFP upon their growth in the non-permissive conditions. Percentages of cells with different pattern of dynein signal are quantitatively represented in the bar graph. The yellow bar represents cells with clustered dynein puncta both in mother and daughter bud in large budded cells. The green bar represents cells with multiple dynein puncta only in the mother bud of large-budded cells, and blue bar shows % cells with no dynein puncta either the mother and daughter bud of large budded cells. The data represent mean ± SD for three independent biological replicates with n ≥ 34 large-budded cells. Bar, 5µm. This experiment comes with a technical limitation to capture the nuclear position with respect to dynein position. The low sample size (metaphase cells) in *slu7kd* overexpressing PAC1 minigene is because high proportion of cells arrested post mitosis. **D.** qRT-PCR to assess the deregulation of cytokinesis-related genes in knockdown and wildtype cells after the shift into non-permissive media for 12 hours.

**Supplementary table I: Strains used in this study**

| Strain name | Genotype | Reference |
|---|---|---|
| H99 | *Wildtype* | (Perfect et al., 1993) |
| Slu7 KD | *MATα Slu7::GAL7p-mCherry-SLU7-HygB* | This study |
| H4 GFP WT | *MATα H99::GFP-H4-NAT* | (Kozubowski et al., 2013) |
| H4 GFP Slu7 KD | *MATα H99::GFP-H4-NAT, SLU7::GAL7p-mCherry-SLU7-HygB* | This study |
| CENPA GFP WT | *MATα H99::GFP-CENP-A-NAT* | (Varshney et al., 2019) |
| CENPA GFP Slu7 KD | *MATα H99::GFP-CENP-A-NAT, SLU7::GAL7p-mCherry-SLU7-HygB* | This study |
| TUB1 GFP WT | *MATα* H99::GFP-tubulin-NAT (pLKB35) | (Yadav and Sanyal, 2018) |
| TUB1 GFP Slu7 KD | *MATα* H99::GFP-tubulin-NAT, *SLU7::GAL7p-mCherry-SLU7-HygB* | This study |
| MAD2Δ H4 GFP WT | *MATα H99 H4::H4-GFP-NAT, mad2Δ::NEO* | (Sridhar et al., 2021) |
| MAD2Δ H4 GFP Slu7 KD | *MATα H99 H4::H4-GFP-NAT, mad2Δ::NEO, SLU7::GAL7p-mCherry-SLU7-HygB* | This study |
| DYN1 3XGFP WT | *MATα DYN1:: DYN1p-DYN1-3xGFP-NEO* | (Chatterjee et al., 2021) |
| DYN1 3XGFP WT Slu7 KD | *MATα DYN1:: DYN1p-DYN1-3xGFP-NEO, SLU7::GAL7p-mCherry-SLU7-HygB* | This study |
| Slu7 FL safe haven | *MATα Slu7::GAL7p-mCherry-SLU7-HygB, SF::Slu7p-Slu7FL-NAT* | This study |
| PAC1 cDNA safe haven GFP H4 | *MATα GFP-H4 Slu7::GAL7p-mCherry-SLU7-HygB, SF::H3p-PAC1 cDNA-NEO* | This study |
| PAC1 cDNA safe haven DYN1 3XGFP | *MATα DYN1 3X GFP Slu7::GAL7p-mCherry-SLU7-HygB, SF::H3p-PAC1 cDNA-NAT* | This study |

**Supplementary table II: Primers used in this study**

| Primer name | Sequence 5' – 3' |
|---|---|
| Slu7 F1 FP | TCA GAG CTC AAT CTC GTA CGC TCA T |
| Slu7 F1 RP | TGAGAGCTCTGTGAAAATGGGTAAGATG |
| Slu7 F2 FP | TTT GTCGAC ATG CTC AGC ACA AG |
| Slu7 F2 RP | TTAGGGCCCGAAGTTTCTGCATG |
| mCh FP | AACATCAAGTTGGACATCACCTCCCA |
| Slu7 +1KB RP | TTATACGTTAGACGATGCCCTGTTG |
| Slu7 FL FP | AACTGCAGCCATCCCTAGGGTA |
| Slu7 FL RP | ACGCGTCGACCATAAGCGGACTT |
| PAC1 cDNA FP | TTACTAGTATGGAGCGCACCCTCAAG |
| PAC1 cDNA RP | TTACTAGTGGGACAGGAATGAACTTGAA |
| SF FP1(Arras et al., 2015) | GGGTATGCCACAGATGCAGAT |
| SF FP2 (Arras et al., 2015) | TCAGCAACGCCGTTGAATCCT |
| SF RP1(Arras et al., 2015) | ACTGGTGAGTACTCAACCAAG |
| SF RP2(Arras et al., 2015) | TTGGATCCTCAATTGTCTCCT |
| PAC1 E1 FP | TCGTCGACGTCGACTTTGAT |
| PAC1 E2 RP | TTATCGCGGCTGGCTGAAACGA |
| PAC1 E2 FP | TCGTTTCAGCCAGCCGCGATAA |
| PAC1 E3 RP | GAACAACCTCTCTGACCCATT |
| PAC1 E3 FP | TGGGTCAGAGAGGTTGTTCCGT |
| PAC1 E4 RP | CCCGTTGAAAAGTCCCATACACG |
| PAC1 E4 FP | CGTGTATGGGACTTTTCAACGGG |
| PAC1 E5 RP | GTAGCTACATAGACACCGGGT |
| PAC1 E5 FP | ACCCGGTGTCTATGTAGCTAC |
| PAC1 E6 RP | CGTTTTCGTGCATCTGCCGTT |
| PAC1 E6 FP | CGTTTTCGTGCATCTGCCGTT |
| PAC1 E7 RP | AATTTTAGAGTCCTGCTAGGGT |

**Supplementary table III: List of DEGs in *slu7kd vs* wildtype**
